# martini: an R package for genome-wide association studies using SNP networks

**DOI:** 10.1101/2021.01.25.428047

**Authors:** Héctor Climente-González, Chloé-Agathe Azencott

**Affiliations:** RIKEN AIP, Tokyo, Japan; MINES ParisTech, PSL Research University, CBIO-Centre for Computational Biology, F-75006 Paris, France; Institut Curie, PSL Research University, F-75005 Paris, France; INSERM, U900, F-75005 Paris, France

**Keywords:** GWAS, networks, R, systems biology, SNP, Bioconductor

## Abstract

Systems biology shows that genes related to the same phenotype are often functionally related. We can take advantage of this to discover new genes that affect a phenotype. However, the natural unit of analysis in genome-wide association studies (GWAS) is not the gene, but the single nucleotide polymorphism, or SNP. We introduce *martini*, an R package to build SNP co-function networks and use them to conduct GWAS. In SNP networks, two SNPs are connected if there is evidence they jointly contribute to the same biological function. By leveraging such information in GWAS, we search SNPs that are not only strongly associated with a phenotype, but also functionally related. This, in turn, boosts discovery and interpretability. *Martini* builds such networks using three sources of information: genomic position, gene annotations, and gene-gene interactions. The resulting SNP networks involve hundreds of thousands of nodes and millions of edges, making their exploration computationally intensive. *Martini* implements two network-guided biomarker discovery algorithms based on graph cuts that can handle such large networks: SConES and SigMod. They both seek a small subset of SNPs with high association scores with the phenotype of interest and densely interconnected in the network. Both algorithms use parameters that control the relative importance of the SNPs’ association scores, the number of SNPs selected, and their interconnection. *Martini* includes a cross-validation procedure to set these parameters automatically. Lastly, *martini* includes tools to visualize the selected SNPs’ network and association properties. *Martini* is available on GitHub (hclimente/martini) and Bioconductor ([martini](https://www.bioconductor.org/packages/release/bioc/html/martini.html)).

## 1 Introduction

Networks are a compact way to integrate information about how genes, proteins and other biomolecules relate to each other. Hence, they frame each measurement from omics experiments within its biological context. We focus here on SNP networks, which model the genome by capturing functional relationships between SNPs. Using such networks in the context of genome-wide association studies (GWAS) boosts discovery of susceptibility SNPs and provides more interpretable hypotheses [3]. In this note, we introduce *martini*, an R package that provides tools to build SNP networks and use them to guide GWAS.

## 2 Main functionalities

In the following sections, we present the different functionalities of *martini* (see Fig 1 for an overview).

**Figure 1:**
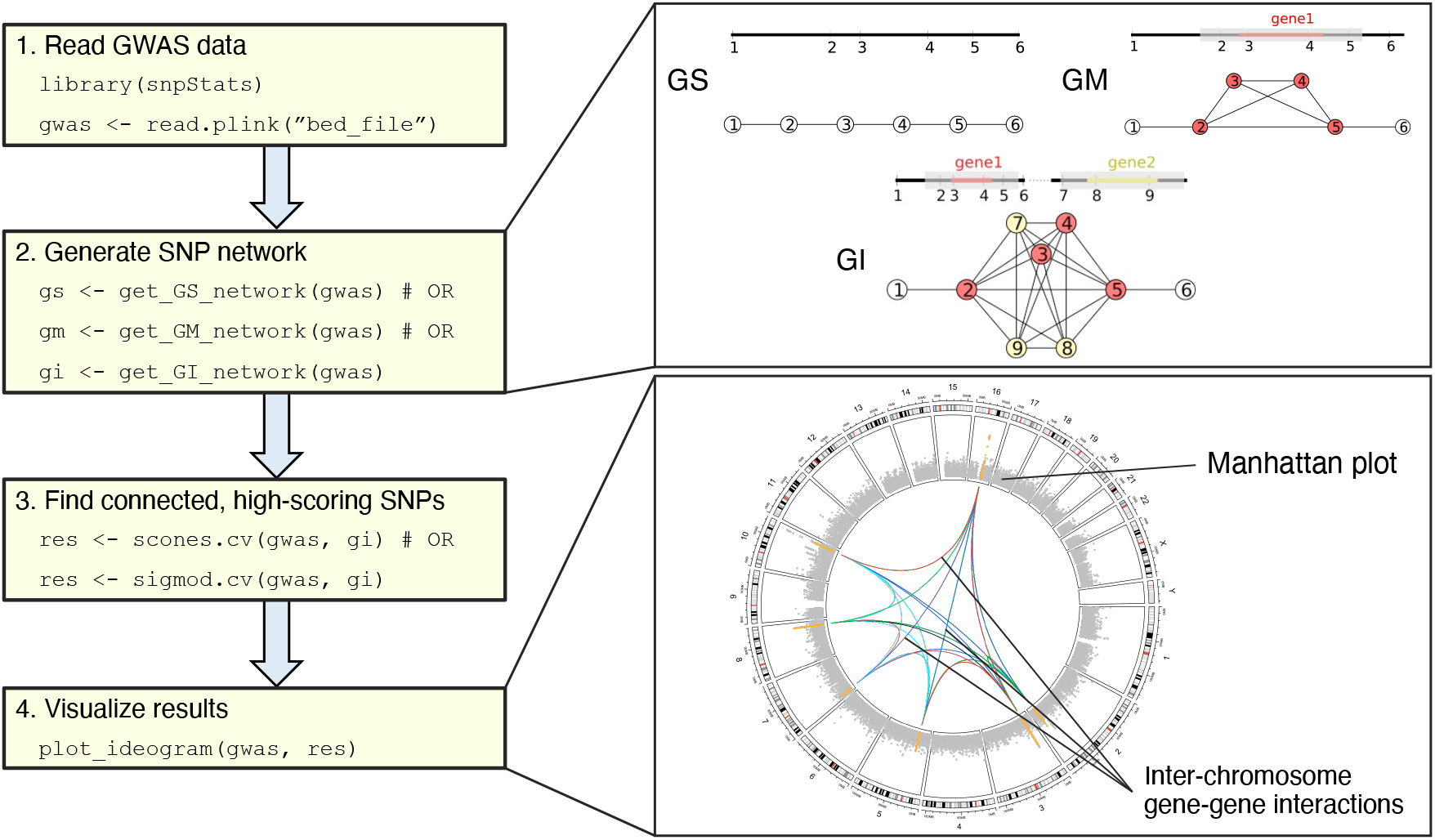
Overview of a the 4 main steps of a *martini* analysis. Adapted from Azencott *et al.* [1].

### 2.1 Building SNP networks

In SNP networks, nodes are SNPs, which are connected by edges when there is some evidence of shared biological function between them. In principle, they can be built from any source of evidence of such shared functionality. *Martini* includes functions to generate the three SNP networks described in Azencott *et al.* [1], so-called GS, GM and GI (Fig 1):

- `get_GS_network()` provides the Genetic Sequence (GS) network, in which SNPs are connected if they are adjacent on the chromosome.
- `get_GM_network()` produces the Gene Membership (GM) network, which includes the GS network and, in addition, interconnects all SNPs that are mapped to the same gene.
- `get_GI_network()` generates the Gene Interaction (GI) network, which includes the GM network and, on top of it, interconnects all the SNPs mapped to two genes that encode interacting proteins.

#### Mapping SNPs to genes

The two latter functions require the user to provide a mapping of SNPs to genes. *martini* provides a convenient way to obtain such a mapping via `snp2ensembl()`, which maps each SNP to all Ensembl genes with overlapping genomic coordinates. This mapping corresponds to the one considered in Azencott *et al.* [1]. In addition, users can easily provide their own mappings to generate other SNP networks. For instance, providing a list of eQTLs together with their target genes (e.g., obtained from GTEx [4]) to `get_GM_network()` will generate networks based on gene expression regulation.

#### Gene-gene interactions

`get_GI_network()` requires the user to provide a list of gene-gene interactions. The `get_gxg()` function of *martini* recovers gene-gene interactions from the BioGRID [6] or STRING [7]. In addition, users can provide their own list of gene-gene interactions beyond protein-protein interactions via a two-column data.frame containing gene-gene pairs.

### 2.2 Finding biomarkers using SConES and SigMod

*Martini* implements two algorithms to find connected subsets of SNPs associated with the phenotype: SConES [1] and SigMod [5]. They share the following formulation:

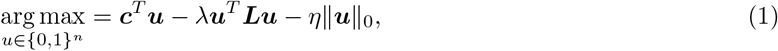

where *u* is a selection vector, in which element *u*_*i*_ is 1 when SNP_*i*_ is selected, and 0 otherwise; ***c*** is a scoring vector, in which element *c*_*i*_ is a measure of association between SNP_*i*_ and the phenotype; *L* is the Laplacian matrix of the SNP network; and *λ* > 0 and *η* > 0 are parameters controlling connectivity and sparsity, respectively. In the case of SConES, *c*_*i*_ = *z*_*i*_, where *z*_*i*_ is a statistical measure of association between SNP_*i*_ and the phenotype. For SigMod, however, *c*_*i*_ = *z*_*i*_ + *λd*_*i*_, where *d*_*i*_ is the number of neighbors SNP_*i*_ in the network. This difference implies that where SConES penalizes the presence of edges connecting selected SNPs with non-selected SNP, SigMod encourages that selected SNPs are connected with each other. In other words, SigMod selects densely connected subnetworks, while SConES selects relatively isolated subnetworks.

#### 2.2.1 Parameter selection

Both SigMod and SConES use two parameters: *λ* and *η*. If both can be provided, *martini*’s functions `scones()` and `sigmod()` provide the corresponding subset of SNPs. However, in most cases, the optimal values of *λ* and *η* are unknown. For such cases, we provide the functions `scones.cv()` and `sigmod.cv()`. These functions explore a grid of parameters in a 10-fold cross-validated setting. A score is computed for each combination of parameters, using the average across the folds of a user-specified scoring function. Then, the best-scoring set of parameters is used in a run on the whole dataset.

*Martini* includes three types of scoring functions: stability, penalized log-likelihood, and network properties. *Stability* selects the parameters that most consistently select the same SNPs across folds. *Penalized log-likelihood* measures are computed on a linear model trained to predict the phenotype using the selected SNPs exclusively. They favor sets of SNPs that lead to good linear predictors but penalize high complexities. *Martini* has three such information criteria available: Bayesian, Akaike, and corrected Akaike (see Appendix A for details). Lastly, *network properties* include two measures that quantify the solution’s edge density: the global and the local clustering coefficients.

Hence, *martini*’s implementation of SigMod is different from the one in the original paper [5], in that we conduct the parameter selection by cross-validation.

#### 2.2.2 Association tests

*Martini* can perform two tests of association between SNPs and the phenotype: 1 d.f. *χ*^2^ and generalized linear models (GLM). The former sets *z*_*i*_ in Eq 1 to the *χ*^2^ test statistic of association between SNP_*i*_ and the phenotype. Hence, it requires the phenotype to be discrete (e.g., case-control). The latter sets *z*_*i*_ to the *χ*^2^ test statistic for the significance of the regression coefficient of SNP_*i*_ in a multivariate GLM explaining the phenotype from SNP_*i*_ as well as additional user-specified covariates, such as principal components to capture population structure. This model can handle both discrete and continuous phenotypes, through the specification of different distribution families. The user can also choose different link functions for the GLM.

### 2.3 Visualization

*Martini*’s `plot_ideogram()` function displays the results on a three-layer ideogram (Fig 1). The first layer displays the cytobands. The second layer contains a circular Manhattan plot showing the statistical association of each SNP with the phenotypes. Non-selected SNPs are colored in gray, and selected SNPs in orange. Lastly, the third layer displays the edges in the SNP network between SNPs from different chromosomes.

## 3 Implementation and availability

*Martini* is implemented in R, and includes a fast C++ implementation of the min-cut/max-flow algorithm [2]. *Martini* is available on GitHub (hclimente/martini) and Bioconductor ([martini](https://www.bioconductor.org/packages/release/bioc/html/martini.html)). The code is licensed as GPL-3. It includes vignettes to show the basic functionalities.

## Funding and acknowledgments

This project was supported by funding from Agence Nationale de la Recherche (ANR-18-CE45-0021-01), the RIKEN Special Postdoctoral Researcher Program, and European Union’s Horizon 2020 research and innovation program (Marie Skłodowska-Curie [666003]). We thank Vivien Goepp for fruitful discussions about network methods and their algorithmic implementations.

## A Log-likelihood penalized scores

Determining optimal values for *λ* and *η* can be seen as a model selection problem and solved using cross-validation, meaning that for each pair of values of these parameters that is considered, one evaluates the cross-validated performance of a penalized logistic regression trained on the features selected by solving Eq 1. However, using the accuracy of this penalized logistic regression as a criterion for evaluation is prone to overfitting. An alternative is to use penalized log-likelihood criteria, which improve generalization by including a regularization term. They take the form

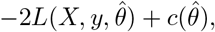

where 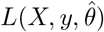 is the log-likelihood of the model, which depends on the design matrix *X*, the outcome vector *y*, and the parameters 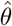; and 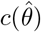 is a measurement of the model’s complexity. We implemented the three most common of these model complexities, resulting in the information criterion (AIC), the Bayesian information criterion (BIC), and the corrected Akaike information criterion (AICc). For all three information criteria, the model complexity is proportional to the number *p*_*in*_ of parameters of the model, here corresponding to the number of selected SNPs:

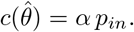

For AIC, the factor *α* is equal to *α* = 2.

For BIC, *α* = ln(*n*) where *n* is the number of samples.

For AICc, which is a modification of AIC proposed for settings where the number of features is much larger than the number of samples, as is the case in GWAS,

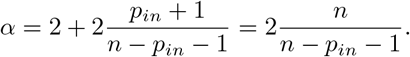

